# IFT-A deficiency in juvenile mice impairs biliary development and exacerbates ADPKD liver disease

**DOI:** 10.1101/2020.09.10.289645

**Authors:** Wei Wang, Tana S Pottorf, Henry H Wang, Ruochen Dong, Matthew A. Kavanaugh, Joseph T. Cornelius, Udayan Apte, Michele T Pritchard, Madhulika Sharma, Pamela V Tran

## Abstract

Polycystic liver disease (PLD) is characterized by the growth of numerous biliary cysts and presents in patients with Autosomal Dominant Polycystic Kidney Disease (ADPKD), causing significant morbidity. Interestingly, deletion of intraflagellar transport-B (IFT-B) genes in adult mouse models of ADPKD attenuates severity of PKD and PLD. Here we examine the role of deletion of IFT-A gene, *Thm1*, in PLD of juvenile and adult *Pkd2* conditional knock-out mice. Perinatal deletion of *Thm1* results in disorganized and expanded biliary regions, biliary fibrosis, shortened primary cilia on CK19+ biliary epithelial cells, and reduced Notch signaling. In contrast, perinatal deletion of *Pkd2* causes PLD, with multiple CK19+ biliary epithelial cell-lined cysts, fibrosis, lengthened primary cilia, and increased Notch and ERK signaling. Perinatal deletion of *Thm1* in *Pkd2* conditional knock-out mice increased hepatomegaly and liver necrosis, indicating enhanced liver disease severity. In contrast to effects in the developing liver, deletion of *Thm1* in adult mice, alone and together with *Pkd2*, did not cause a biliary phenotype nor affect *Pkd2*-mutant PLD, respectively. However, similar to juvenile PLD, Notch and ERK signaling were increased in adult *Pkd2*-mutant cyst-lining cholangiocytes. Taken together, *Thm1* is required for biliary tract development, likely by enabling Notch signaling, and proper biliary development restricts PLD severity. Unlike IFT-B genes, *Thm1* does not affect hepatic cystogenesis, suggesting divergent regulation of signaling and cystogenic processes in the liver by IFT-B and –A. Notably, increased Notch signaling in cyst-lining cholangiocytes may indicate that aberrant activation of this pathway promotes hepatic cystogenesis, presenting as a novel potential therapeutic target.

## Introduction

Hepatorenal fibrocystic diseases comprise a spectrum of diseases characterized by varying degrees of cysts and fibrosis in the kidney and liver. These diseases include Autosomal Dominant Polycystic Kidney Disease (ADPKD), which is among the most common, life-threatening monogenetic diseases, affecting 1:500 individuals worldwide. In addition to developing renal cystic disease, up to 94% of individuals with ADPKD manifest Polycystic Liver Disease (PLD), which contributes significant morbidity. In PLD, numerous fluid-filled cysts derive from the biliary ducts and progressively burden the liver, causing hepatomegaly, gastrointestinal and respiratory discomfort, and pain. Infection and hemorrhaging of cysts can also arise^1,2^. While Tolvaptan, the only FDA-approved therapy for ADPKD^3,4^, attenuates the renal cystic disease, reports of its effects in the liver conflict. A case report showed reduced liver volume in an ADPKD patient^5^, but several studies indicate Tolvaptan can cause liver toxicity^4,6-8^. The paucity of therapies and conflicting reports reflect the need to continue uncovering mechanisms underlying liver fibrocystic disease.

ADPKD is caused by mutations in *PKD1* or *PKD2*, encoding polycystin proteins, which form a complex and function at the primary cilium. Mutation of *PKD1* or *PKD2* typically results in deficiency of the polycystin complex in primary cilia^9,10^. These antenna-like organelles sense chemical and mechanical cues in the extracellular environment and mediate signaling pathways. In the liver, primary cilia project from the apical membrane of cholangiocytes into the biliary lumen, where they sense bile composition and flow. Primary cilia are dynamic structures built and maintained by intraflagellar transport (IFT), which mediates the bi-directional transport of protein cargo along the ciliary microtubular axoneme. IFT complex B (IFT-B) interacts with the kinesin motor to mediate anterograde IFT, while IFT-A together with cytoplasmic dynein is required for retrograde IFT. IFT-A also mediates ciliary entry of membrane and signaling molecules^11^. Reflecting their differential roles in ciliogenesis and maintenance, deletion of an IFT-B gene usually results in absence of cilia^12^, while deletion of an IFT-A gene causes shortened cilia with accumulation of proteins in a bulbous distal tip^13^. Deletion of IFT-B versus IFT-A genes can also result in differential regulation of signaling pathways.

Mutations of the IFT-B gene, *IFT56*, have been reported in families with biliary ciliopathies^14^. In mice, mutation of IFT-B gene, *Ift88*, causes rapid expansion of the biliary regions with an increased number of cholangiocytes and periportal fibrosis^15^. Interestingly, deletion of IFT-B genes, *Kif3a* or *Ift20*, in adult *Pkd1* conditional knock-out (cko) mice, and of IFT-B gene, *Ift88* in adult *Pkd2* cko mice, attenuates both renal and biliary cystogenesis^16,17^, suggesting the ciliome has therapeutic potential. The mechanisms by which cilia loss or mutation attenuates ADPKD cystogenesis remain largely undefined. Recently, P53 and Wnt signaling have been reported as potential pathways that are suppressed by cilia mutation in *Pkd; Ift-B* mutant renal epithelial cells^17,18^. However, signaling pathways in the liver of *Pkd; Ift* double mutant mice have not been studied. Additionally, the role of IFT-A in ADPKD liver disease is unknown. To elucidate a broader mechanism of ciliopathic liver disease, we investigate the role of murine IFT-A gene, *Thm1* (also known as *Ttc21b*), in developing and mature liver, alone and in conjunction with deletion of *Pkd2. THM1* mutations have been identified to cause nephronophthisis and to modify severity of Bardet Biedl Syndrome, Jeune Syndrome, and Meckel Syndrome, ciliopathies which manifest fibrocystic liver disease^19^. We have shown that deletion of *Thm1* results in various ciliopathy phenotypes, including developmental defects^13^, renal cystic disease^20^ and obesity^21^. Here we show that *Thm1* is required for postnatal biliary tract development, as well as Notch signaling. Further, we identify increased Notch signaling in cyst-lining cholangiocytes, suggesting aberrant activation of Notch signaling as a potential contributor to hepatic cystogenesis.

## Methods

### Mice

*Pkd2*^*flox/flox*^ and *ROSA26-Cre* mice were obtained from the Jackson Laboratories (Stock numbers 017292 and 004847, respectively). Generation of *Thm1* cko mice has been previously described ^20^: *Thm1*^*aln/+*^; *ROSA26Cre*^*ERT+*^ male mice were mated with *Thm1*^*flox/flox*^ females. *ROSA26-Cre*^*ERT/+*^ and *Pkd2* ^*flox*^ alleles were introduced into the colony to generate *Thm1*^*flox/flox*^;*Pkd2*^*flox/flox*^ or *Thm1*^*flox/flox*^;*Pkd2*^*flox/+*^ females and *Pkd2*^*flox/flox*^; *Thm1*^*aln/+*^, *ROSA26-Cre*^*ERT/+*^ males, which subsequently were crossed. To generate early-onset models of ADPKD, nursing mothers were injected intraperitoneally at postnatal day 0 (P0) with tamoxifen (8mg/40g; Sigma) to induce gene deletion. Mice were sacrificed at P21. To generate late-onset models, offspring of matings between *Thm1*^*flox/flox*^;*Pkd2*^*flox/flox*^ or *Thm1*^*flox/flox*^;*Pkd2*^*flox/+*^ females with *Pkd2*^*flox/flox*^; *Thm1*^*aln/+*^, *ROSA26-Cre*^*ERT/+*^ males were injected intraperitoneally at P28 with tamoxifen (8mg/40g). Mice were sacrificed at 6 months of age. All mouse lines were maintained on a pure C57BL6/J background (backcrossed 10 generations).

### Liver and body weight measurements

Livers were dissected and weighed using a standard laboratory weighing scale. Liver weight/body weight (LW/BW) ratios were calculated as liver weight divided by body weight for each mouse.

### Histology

The left lobe of the liver was fixed in 10% formalin for several days and then processed in a tissue processor and embedded in paraffin. Tissue sections (7µm) were obtained with a microtome. Sections were deparaffinized, rehydrated through a series of ethanol washes and stained with hematoxylin and eosin (H&E) or picrosirius red according to standard protocols^22^. Images were taken with a Nikon 80i microscope equipped with a Nikon DS-Fi1 camera. ImageJ was used to quantify cystic areas and necrotic areas in H&E-stained sections. Percent cystic index was calculated by dividing the area of liver cysts over the area of the whole liver section, then multiplying by 100%. Percent necrosis was calculated by dividing the area of liver necrotic sites over the area of the whole liver section, then multiplying by 100%.

### Immunofluorescence

Following deparaffinization and rehydration described above, tissue sections were subjected to antigen retrieval. Tissue sections were steamed for 15 minutes in Sodium Citrate Buffer (10 mM Sodium Citrate, 0.05% Tween 20, pH 6.0), returned to room temperature, rinsed 10 times in distilled water, washed 5 minutes in PBS, incubated for 5 minutes in 1% SDS in PBS based on a method by Brown et al., 1996^23^, then washed 3 times in PBS. Sections were blocked with 1% BSA in PBS for 1 hour at room temperature, and then incubated with primary antibodies against acetylated α-tubulin (1:4000; Sigma), αSMA (1:500; Abcam), PCNA (1:300; Cell Signaling), CK19 (1:200; Abcam), F4/80 (1:400; Cell Signaling) and P-ERK (1:200; Cell Signaling) overnight at 4°C. Sections were washed three times in PBS, and then incubated with secondary antibodies conjugated to Alexa Fluor 488 or Alexa Fluor 594 (1:500; Invitrogen by Thermo Fisher Scientific) for 1 hour at room temperature. After three washes of PBS, sections were mounted with Fluoromount-G containing 4′,6-diamidino-2-phenylindole (DAPI) (Electron Microscopy Sciences). Staining was visualized and imaged using a Nikon 80i microscope with a Nikon DS-Fi1 camera or a Nikon Eclipse TiE attached to an A1R-SHR confocal, with an A1-DU4 detector, and LU4 laser launch.

### Measurement of Mean Fluorescence Intensity

Image J was used to measure Mean Fluorescence Intensity (MFI). Integrated density of P-ERK^+^ cyst-lining epithelia was measured. Five regions adjacent to P-ERK^+^ cyst-lining epithelia with low fluorescence were selected to calculate mean background fluorescence. MFI was calculated by dividing integrated density of selected cyst-lining epithelial cells over the area of selected cyst-lining epithelial cells, then subtracting the mean background fluorescence.

### Cilia length quantification

Tissue sections immunostained for CK19 and acetylated α-tubulin were imaged at multiple planes. Cilia images were converted to black and white. Cilia lengths were quantified using ImageJ.

### Immunohistochemistry

Following deparaffinization and rehydration described above, tissue sections were subjected to antigen retrieval, performed by steaming sections for 25 minutes in Sodium Citrate Buffer (10 mM Sodium Citrate, 0.05% Tween 20, pH 6.0). To minimize background staining, sections were treated with 3% hydrogen peroxide for 30 min, washed in PBS, then blocked with 1% BSA for 1 hour. Tissue sections were incubated with primary antibodies against Jagged 1 (1:50; Abcam); Notch 2 (1:100; LSBio) and Presenillin-1 (1:100; Genscript) overnight at 4°C. Following 3 washes in PBS, sections were incubated with HRP-conjugated rabbit secondary antibody (Cell Signaling) for 30 minutes. Following another 3 washes in PBS, tissues were incubated with ABC reagent (Vector Laboratories), rinsed in PBS, and then incubated with SigmaFAST DAB metal enhancer (Sigma) until desired signal/color was obtained, then counterstained with haemotoxylin. Staining was visualized and imaged using a Nikon 80i microscope with a Nikon DS-Fi1 camera.

### Statistics

Statistical significance (P < 0.05) was determined using one-way ANOVA followed by Tukey’s test or unpaired t-test for comparison of more than two groups or of two groups, respectively. GraphPad Prism 8 software was used to perform these analyses.

## Results

### Thm1 loss impairs biliary development and exacerbates severity of polycystic liver disease in juvenile Pkd2 conditional knock-out mice

To explore the role of IFT-A deficiency in the developing liver, we deleted *Thm1* alone, and together with *Pkd2* in mice at postnatal day (P) 0, and examined the liver phenotypes of control, *Thm1* cko, *Pkd2* cko, and *Pkd2*;*Thm1* dko mice at P21. Histology of *Thm1* cko livers revealed expanded and disorganized biliary regions (Figure 1A), without affecting liver and body weights (Figures 1B and 1C)^24^. In contrast, perinatal deletion of *Pkd2* caused multiple liver cysts and increased liver weight to body weight (LW/BW) ratios (Figure 1D). Additional deletion of *Thm1* in *Pkd2* cko mice did not alter hepatic cystogenesis, but increased LW/BW ratios and multifocal necrosis, suggesting increased severity of the liver disease (Figures 1A-1F).

**Figure 1.**
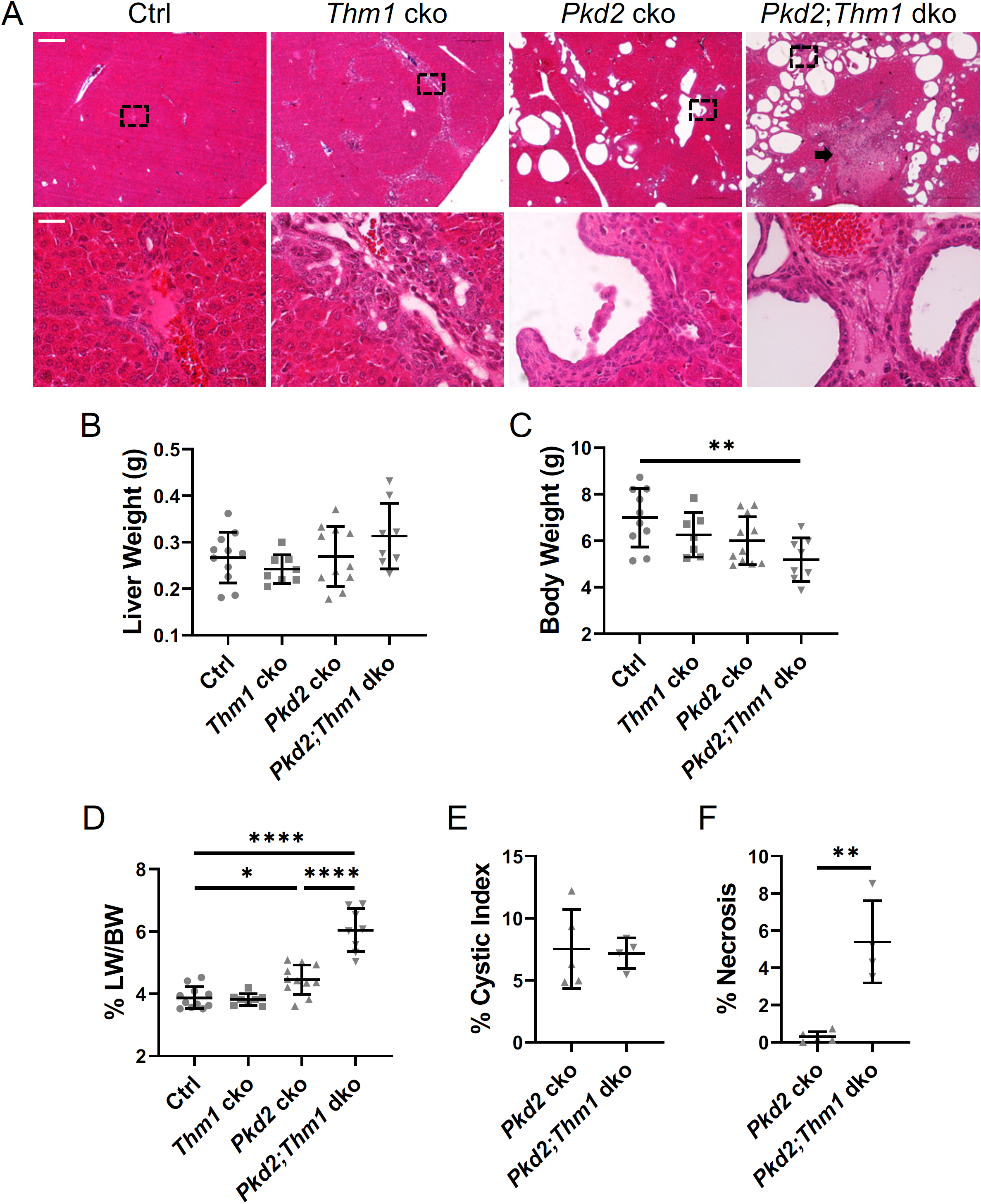
Biliary defects and liver cysts of juvenile *Thm1* cko, *Pkd2* cko, and *Pkd2*;*Thm1* dko mice. (A) H&E-stained liver sections of P21 mice. Dotted boxed regions in upper panels are shown at higher magnification in bottom panels. Scale bars -250 µm (upper); 25 µm (bottom). Arrow points to area of necrosis. N≥4 mice/genotype. (B) Liver weight (C) Body weight (D) Liver weight/body weight (LW/BW) ratios. (E) Liver cystic index (F) Percentage of necrosis. Statistical significance was determined by ANOVA followed by Tukey’s test in (C and D) and by unpaired t-test in (F). *P<0.05; **P<0.01; ****P< 0.0001

Since cysts in PLD originate from bile ducts, we immunostained liver sections for cytokeratin 19 (CK19), a marker of biliary epithelial cells. In *Thm1* cko livers, there was an increased number and a disorganization of CK19+ cells (Figure 2A). Not all cells within the *Thm1* cko expanded biliary regions were CK19+, indicating other cell types also account for the expansion. In *Pkd2* cko and *Pkd2;Thm1* dko mice, all cyst-lining cells were CK19+, consistent with the cysts arising from biliary ducts.

**Figure 2.**
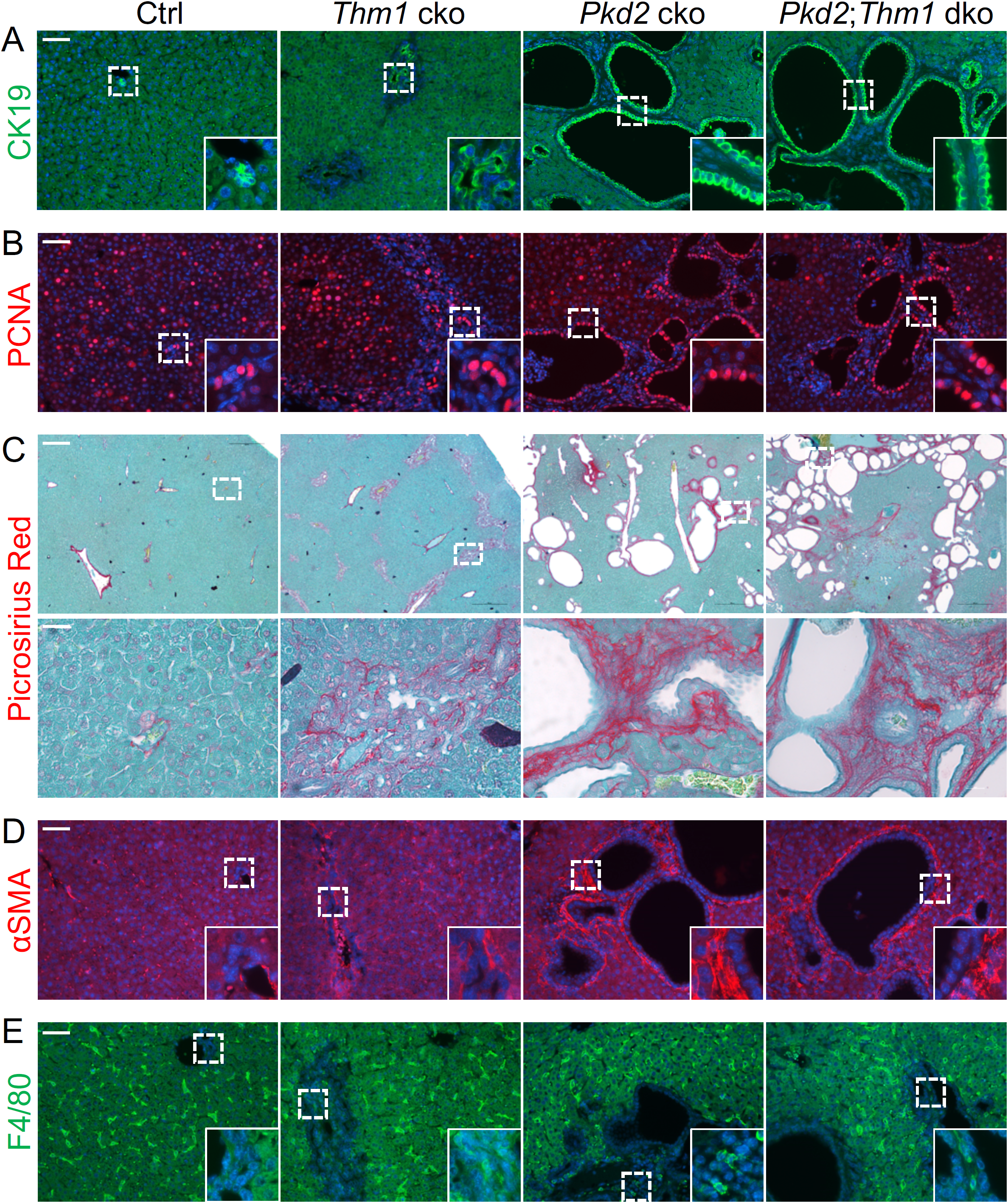
Juvenile *Thm1* cko, *Pkd2* cko and *Pkd2*;*Thm1* dko mice develop liver fibrosis. Immunostaining for (A) CK19+ (green) and (B) PCNA (red). Scale bar -50 µm. Dotted boxed regions in upper panels are shown at higher magnification in insets. (C) Picrosirius red-stained liver sections. Dotted boxed regions in upper panels are shown at higher magnification in bottom panels. Scale bars -250 µm (top); 25 µm (bottom). (D) Immunostaining for αSMA (red) and (E) F4/80 (green). Scale bar -50 µm. Dotted boxed regions in upper panels are shown at higher magnification in insets. N=3 mice/genotype

We next examined proliferation, a cellular hallmark of cystogenesis, by immunostaining for proliferating cell nuclear antigen (PCNA). In mice, livers continue to mature until around 4 weeks of age. At P21, control livers revealed the presence of some PCNA+ hepatocytes and cholangiocytes (Figure 2B). Similarly, *Thm1* cko livers showed some PCNA+ hepatocytes and cholangiocytes. In contrast, in *Pkd2* cko and *Pkd2:Thm1* dko livers, some hepatocytes and many cyst-lining cells were PCNA+, consistent with the notion that proliferation drives cystogenesis.

### Thm1 loss causes biliary fibrosis

Liver fibrosis is a common feature of PLD, presenting in more than half of PLD patients^25^. To assess fibrosis, we stained liver sections with picrosirius red, which labels collagen fibers. In control sections, picrosirius red was present mainly around the blood vessels (Figure 2C). However, in *Thm1* cko livers, picrosirius red staining was present also in the biliary regions. In *Pkd2* cko livers, even more intense staining of picrosirius red was observed surrounding hepatic cysts. A similar level of intense staining was observed in *Pkd2;Thm1* dko livers. We next immunostained for alpha-Smooth muscle actin (αSMA), which labels myofibroblasts, the primary cell types that secrete extracellular matrix during fibrosis. While αSMA was mostly detected around blood vessels in control livers, αSMA was also present in the biliary regions of *Thm1* cko livers and surrounding the cysts of *Pkd2* cko and *Thm1;Pkd2* dko livers (Figure 2D). Myofibroblasts are activated by M1 macrophages, which contribute to pro-inflammatory processes. To assess presence of M1 macrophages, we stained liver sections for F4/80. While F4/80+ cells were present in the sinusoids across all genotypes, F4/80+ cells were present also in the biliary regions of *Thm1* cko, *Pkd2* cko and *Pkd2;Thm1* dko livers (Figure 2E). These data suggest that *Thm1* loss causes biliary fibrosis. While fibrosis also accompanies early onset PLD, additional loss of *Thm1* did not exacerbate PLD fibrosis at P21.

### Cholangiocyte cilia are shortened in Thm1 cko livers, but lengthened in Pkd2 cko livers of juvenile mice

While primary cilia are present on bipotential hepatoblasts, cilia are absent on differentiated hepatocytes, but present on cholangiocytes protruding into the bile ducts^26^. We examined cholangiocyte cilia lengths by co-immunostaining liver sections for acetylated α-tubulin, a marker of the ciliary axoneme, together with either CK19 or with IFT81, a component of the IFT-B complex. *Thm1* cko cholangiocyte cilia were shortened, in contrast to *Pkd2* cko cilia, which were lengthened (Figures 3A and 3B). *Pkd2;Thm1* dko cholangiocyte cilia lengths were shortened and similar to those of *Thm1* cko cilia with some showing accumulation of IFT81 at the distal tip, consistent with a retrograde IFT defect^13^. These data suggest that deletion of *Thm1* in *Pkd2* cko mice controls ciliary length in cholangiocytes.

**Figure 3.**
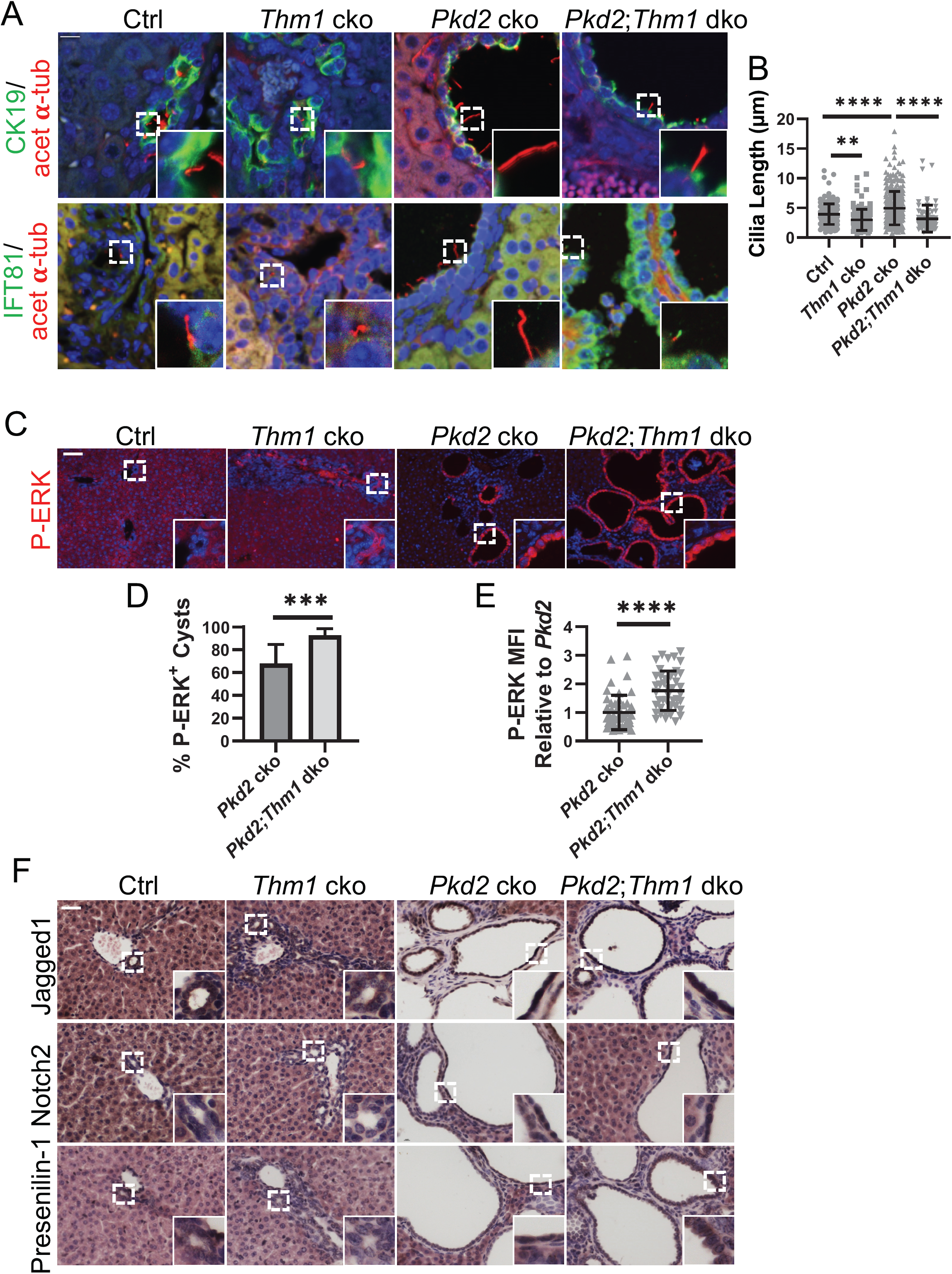
Primary cilia and ERK and Notch signaling of juvenile *Thm1* cko, *Pkd2* cko and *Pkd2;Thm1* dko CK19+ cells. (A) Immunostaining for acetylated α-tubulin (red) and CK19 (green) and for (B) acetylated α-tubulin (red) and IFT81 (green) of P21 mice. Scale bar -10 µm. (C) Quantification of cholangiocyte cilia lengths from (A). Each data point represents an individual cilium from 3 mice/genotype. Statistical significance was determined by ANOVA followed by Tukey’s test. ****P< 0.0001. (D) Immunostaining for P-ERK. Dotted boxed regions in upper panels are shown at higher magnification in insets. N=3 mice/genotype. Scale bar -50 µm. (E) Percent of cysts lined with P-ERK+ epithelia. Three different regions/mouse section were imaged and 115-250 cysts were quantified for each animal. (F) Mean Fluorescence Intensity (MFI) of P-ERK staining. Each dot represents a cyst lined with P-ERK+ epithelia. N=3 mice/genotype. Statistical significance was determined by unpaired t-test. ****P< 0.0005 (G) Immunohistochemistry for Notch signaling components – Jagged1 ligand, Notch2 receptor and Presenilin-1. Dotted boxed regions in upper panels are shown at higher magnification in insets. N=3 mice/genotype

### Notch signaling is downregulated in Thm1 cko biliary regions, while ERK and Notch signaling are increased in Pkd2 cko and Pkd2;Thm1 dko cyst-lining cholangiocytes

Primary cilia mediate signaling pathways. Thus we next examined signaling pathways that are known to play a role in PLD or in biliary development. In PLD, ERK activation in cyst-lining cholangiocytes is a major driver of cell proliferation^27^. To examine the effect of *Thm1* on this pathway in cholangiocytes, we immunostained liver sections for P-ERK. In control livers and similarly, in *Thm1* cko livers, less than half of biliary epithelial cells (approximately 40%) were positive for P-ERK (Figure 3C). However, in *Pkd2* cko and *Pkd2;Thm1* dko livers, approximately 62% and 90%, respectively, of cyst-lining cholangiocytes were P-ERK+ (Figures 3C-3E). Additionally, intensity of P-ERK staining was increased in *Pkd2;Thm1* dko cyst-lining cells relative to *Pkd2* cko. Thus, loss of *Thm1* on a *Pkd2* cko background increases ERK activation in biliary epithelial cells.

The Notch pathway is essential for biliary tract development^28^ and has been shown to be regulated by primary cilia^29^. Thus we examined Notch signaling status in the mutant livers. The Notch cascade initiates with Delta or Jagged ligands that are received by Notch receptors. Of the 4 Notch receptors, Notch 2 is required for cholangiocyte differentiation and tubulogenesis. Notch receptors are cleaved by presenilin-1, forming the Notch Intracellular Domain (NICD). NICD translocates to the nucleus to activate a transcription complex containing Recombination Signal Binding Protein For Immunoglobulin Kappa J Region (RBPj), which drives transcription of target genes. In *Thm1* cko biliary regions, Jagged1, Notch2, and Presenilin-1 were downregulated (Figure 3F). In contrast, in *Pkd2* cko and *Pkd2;Thm1* dko cyst-lining cholangiocytes, Jagged1, Notch2, and Presenilin-1 were increased, suggesting that Notch signaling may promote hepatic cystogenesis.

### Deletion of Thm1 in a mature liver does not affect biliary morphology nor ADPKD hepatic cystogenesis

Since *Ift*-*B* deficiency in adult *Pkd1* or *Pkd2* cko mice attenuates hepatic cystogenesis^16,17^, we next examined the role of IFT-A deficiency in adult, slowly progressive PLD mouse models. We deleted *Thm1* together with *Pkd2* at P28, when livers have matured, and examined control, *Thm1* cko, *Pkd2* cko and *Pkd2;Thm1* dko livers at 6 months of age. Global deletion of *Thm1* once the liver has matured did not affect biliary morphology nor liver weights (Figure S1), but increased body weight, consistent with its role in obesity^21^. In *Pkd2* cko mice, hepatic cysts, hepatomegaly and increased LW/BW ratios were evident (Figures 4A-4D). While ADPKD female patients are associated with more severe PLD^30^, LW/BW ratios were similar between male and female *Pkd2* cko mice. Loss of *Thm1* together with *Pkd2* increased liver weights in female mice, but not in male mice. However, in both females and males, hepatic cystogenesis was similar between *Pkd2* cko and *Pkd2;Thm1* dko mice (Figure 4E), suggesting that *Thm1* deletion did not affect cyst formation and progression *per se*.

**Figure 4.**
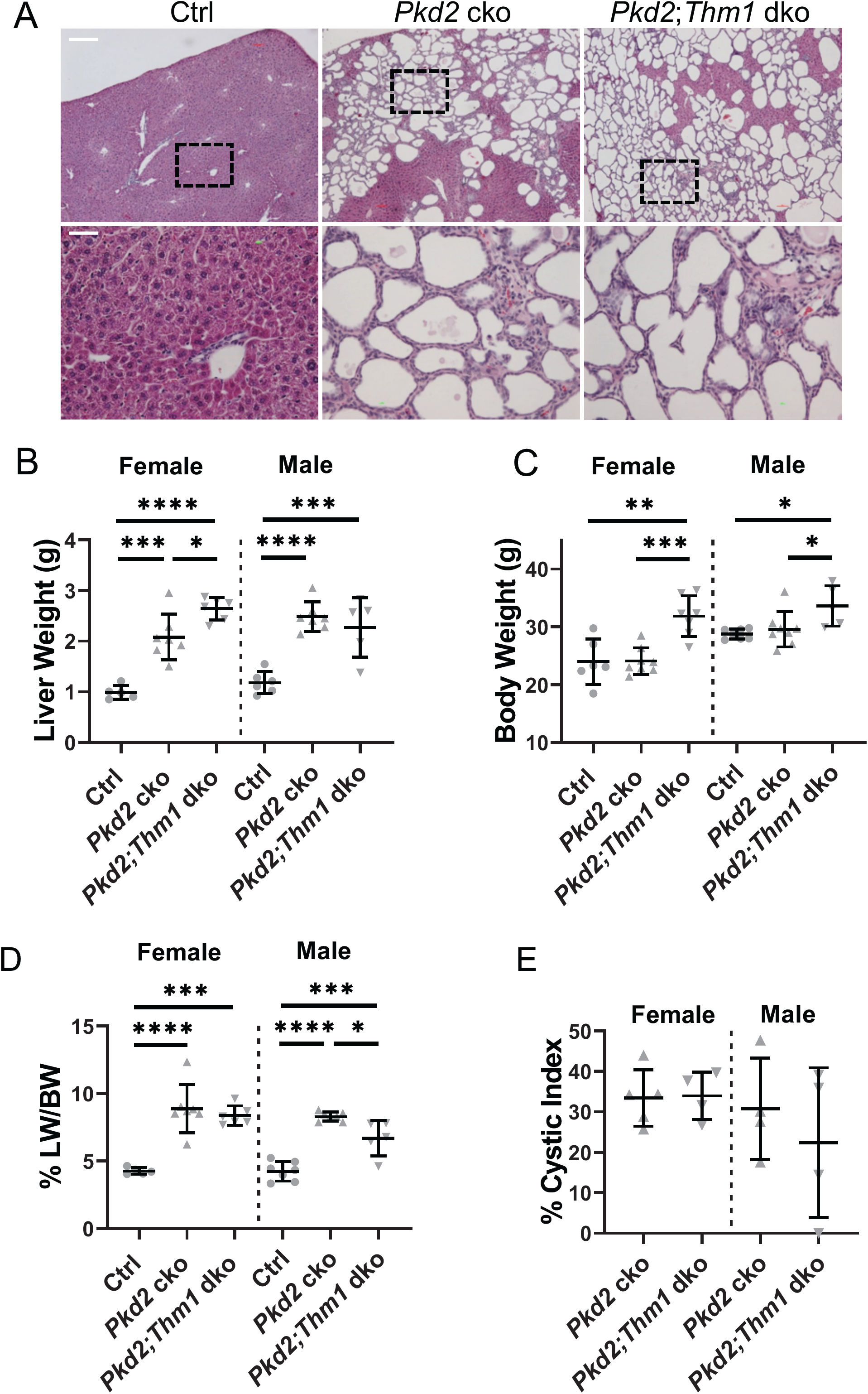
Adult onset loss of *Thm1* does not affect severity of polycystic liver disease in *Pkd2* cko mice. (A) H&E-stained liver sections from male mice at low magnification (top panels). Dotted boxed regions are shown at high magnification (bottom panels). Scale bar -250 µm (top); 25 µm (bottom). (B) Liver weight (B) body weight and (C) and liver weight/body weight ratios of 6-month-old male mice. (D) Liver weight (E) body weight and (F) and liver weight/body weight ratios of 6-month-old female mice Statistical significance was determined by ANOVA followed by Tukey’s test. *P<0.05; **P<0.01; ***P< 0.001; ****P< 0.0001

All cyst-lining cells of adult *Pkd2* cko and *Pkd2;Thm1* dko livers were CK19+ (Figure 5A), indicating their biliary origin. Many cyst-lining cells were also positive for PCNA (Figure 5B), consistent with proliferation being a major contributor of cystogenesis. F4/80+ and alphaSMA+ cells were also present surrounding the biliary cysts (Figures 5C and 5D), indicating the presence of macrophages and myofibroblasts, respectively, and thus, pro-fibrotic processes. In *Thm1* cko livers, immunostaining for CK19 revealed bile ducts similar to control (Figure S2A). There was also an absence of PCNA+ and F4/80+ cells in the *Thm1* cko bile ducts, similar to control (Figures S2B and S2C), substantiating that loss of *Thm1* in a mature liver does not cause a biliary phenotype.

**Figure 5.**
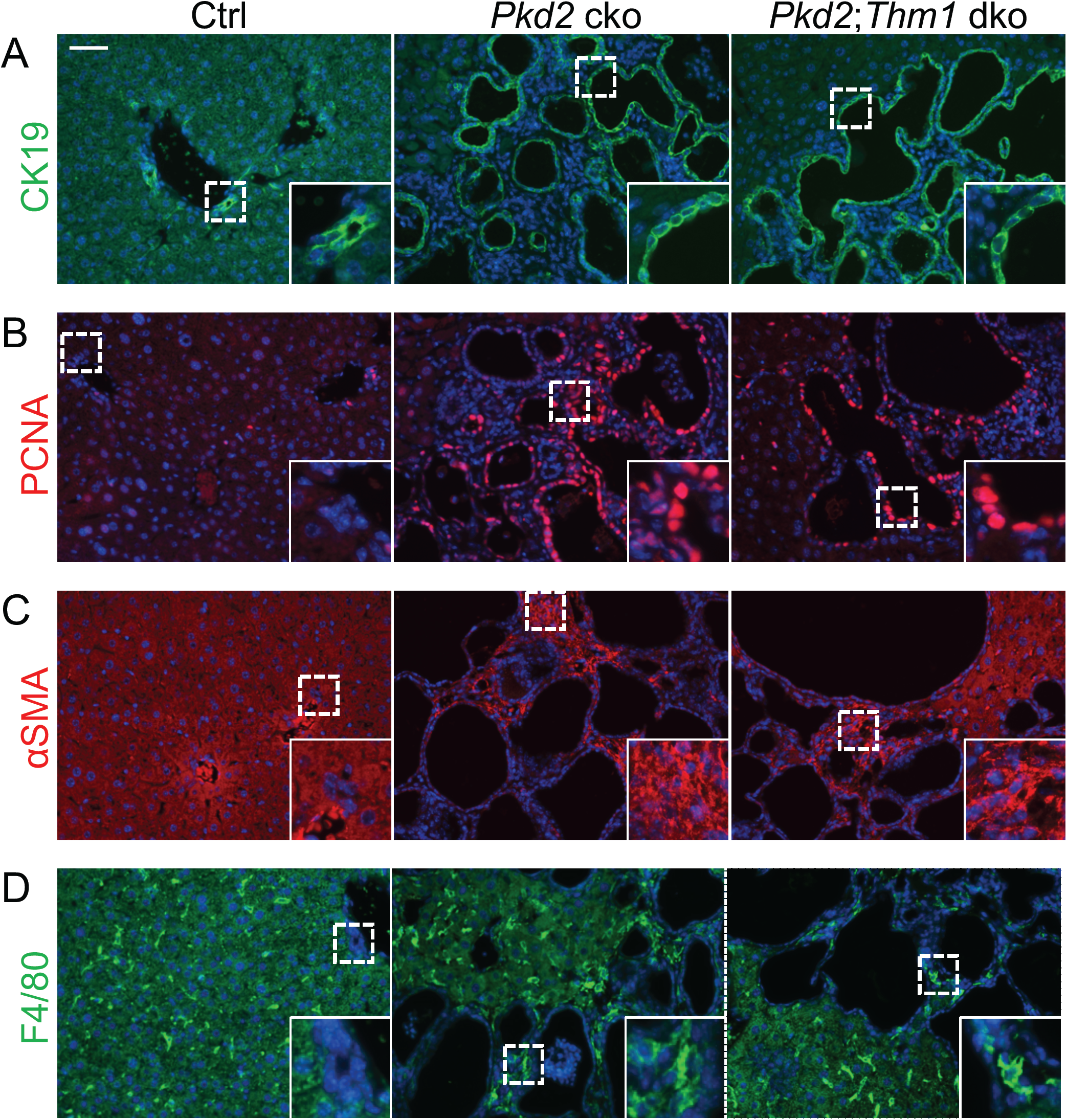
PLD of adult *Pkd2* cko and *Pkd2*;*Thm1* dko mice. Immunostaining for (A) CK19+ (green); (B) PCNA (red); (C) αSMA (red) and (D) F4/80 (green). Dotted boxed regions are shown at higher magnification in insets. Scale bar -50 µm. N=3 mice/genotype

Primary cilia of CK19+ cells were shortened in *Thm1* cko mice (Figures S3A and S3B), suggesting that deletion of *Thm1* did induce a ciliary defect. In *Pkd2* cko mice, cholangiocyte primary cilia showed a wider range of cilia lengths with some reaching over 9μm, while the maximum cilia length in control cholangiocytes was less than 6μm (Figures 6A and 6B). Yet, collectively, cilia lengths of *Pkd2* cko mice were not significantly longer than those of control mice. *Pkd2;Thm1* dko cholangiocyte cilia lengths were shortened relative to those of control and *Pkd2* cko mice, suggesting that loss of *Thm1* influenced the ciliary phenotype.

**Figure 6.**
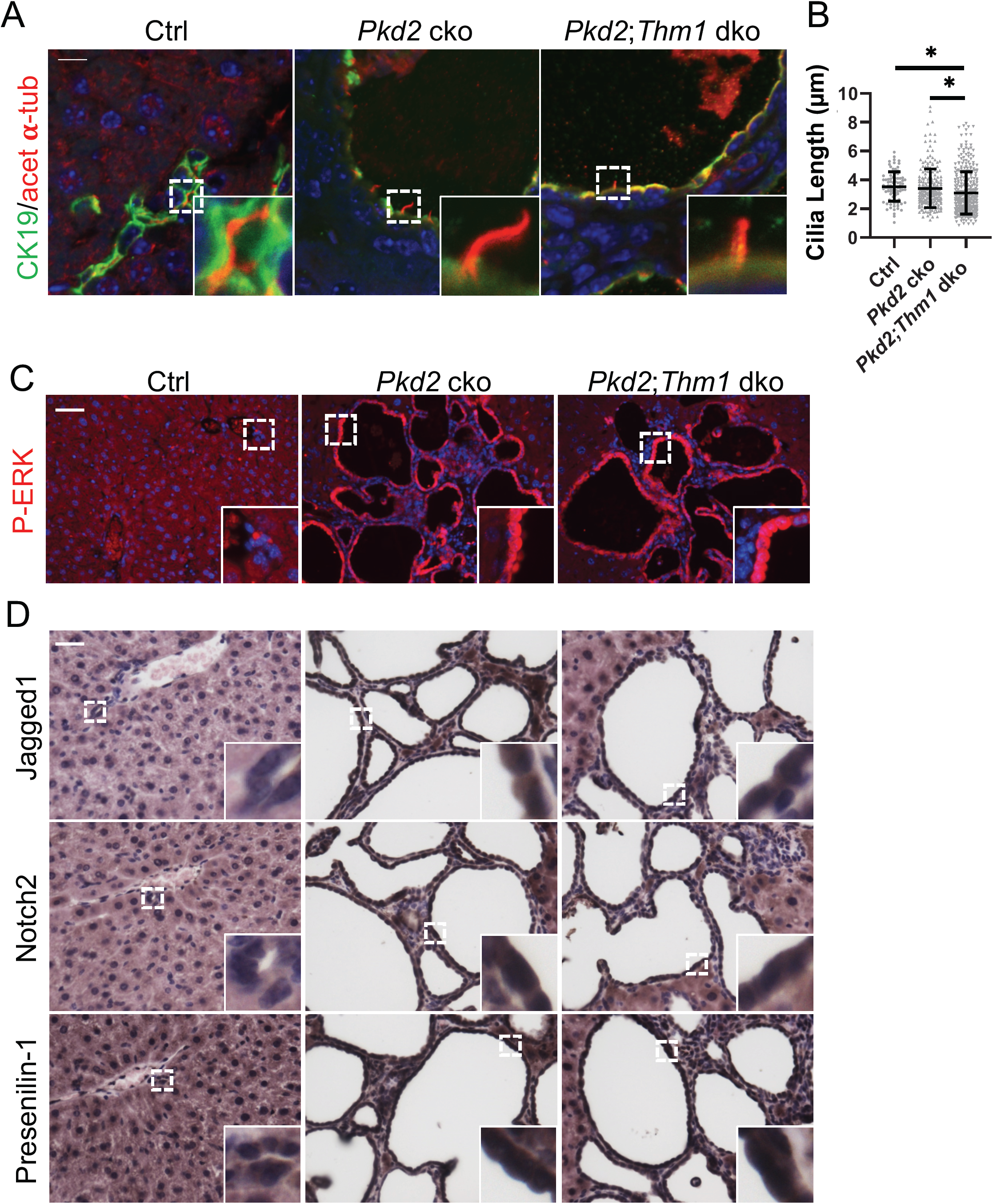
Primary cilia and ERK and Notch signaling of adult *Pkd2* cko and *Pkd2;Thm1* dko CK19+ cells. (A) Immunostaining for acetylated α-tubulin (red) and CK19 (green) of 6-month-old mice. Scale bar -10 µm. (B) Quantification of cilia lengths. Each data point represents an individual cilium from 4 mice/genotype. Statistical significance was determined by ANOVA followed by Tukey’s test. *P< 0.05. (C) Immunostaining for P-ERK. Dotted boxed regions in upper panels are shown at higher magnification in insets. N=3 mice/genotype. Scale bar -50 µm. (D) Immunohistochemistry for Jagged1 ligand, Notch2 receptor and Presenilin-1. Dotted boxed regions in upper panels are shown at higher magnification in insets. N=3 mice/genotype

Finally, while *Thm1* cko bile ducts showed absence of P-ERK and similar intensities of Notch signaling components to those in control bile ducts (Figure S3C), cyst-lining cholangiocytes of *Pkd2* cko and *Pkd2;Thm1* dko bile ducts showed increased P-ERK and increased expression of Jagged1, Notch2, and Presenilin-1 (Figure 8). These data suggest that similar signaling pathways drive PLD in juvenile and adult *Pkd2* cko mice.

## Discussion

During biliary tract development, ductal plates around the portal mesenchyme duplicate, and subsequently form tubules and elongate. These tubular structures form intrahepatic bile ducts (IHBD), which subsequently connect to the extrahepatic hilar duct. Cells that do not integrate into a tubular structure or IHBD, or do not connect with the hilar duct are either apoptosed or lose their cholangiocyte specification and convert into hepatocytes. When these developmental events fail to occur, these unintegrated cells may contribute to ductal plate malformations and result in fibrocystic liver diseases^25,31^. In juvenile *Thm1* cko mice, we observed an increased number of CK19+ cells that were disorganized, as well as biliary fibrosis. Notably, this phenocopies the biliary phenotype of *Notch2* ko mice, namely increased and disorganized pan-cytokeratin-expressing cells surrounded by fibrosis^32^. Additionally, *Notch2* ko pan-cytokeratin+ cells express CK19 less strongly than control biliary epithelial cells. Since CK19+ intensity increases with maturation of the biliary epithelial cell, this suggests *Notch2* ko cells are less differentiated. 3D morphological analysis of the biliary tree in *Notch1/2* ko and in *RBPj* ko livers show a paucity of biliary ducts, suggesting that the expanded biliary regions observed via histology result from lack of apoptosis or regression of undifferentiated cells and unconnected ducts^28^. Similarly, the expanded biliary region in *Thm1* cko livers may result from undifferentiated ductal plates that have failed to apoptose or regress. By extension, the number of functional biliary ducts are likely decreased in *Thm1* cko mice. We noted a downregulation of Notch signaling in *Thm1* cko biliary regions. Since the *Thm1* cko biliary phenotype phenocopies that of *Notch2* ko mice, the downregulation of Notch signaling in *Thm1* cko livers likely causes the impaired biliary development.

In the developing epidermis, Notch signaling has been shown to be mediated by primary cilia. Deletion of *Kif3a* in epidermal cells during embryogenesis results in an absence of primary cilia and downregulation of the Notch pathway^29^. Similarly, reduced Notch signaling in *Thm1* cko biliary epithelial cells may be a direct effect of the shortened cilia defect. In contrast, *Pkd2* mutant primary cilia were lengthened and showed aberrant activation of Notch signaling. On *Pkd1* and *Pkd2*-mutant renal epithelial cells, primary cilia have been shown to be lengthened^17,33,34^, and a recent report suggests that increased primary cilia lengths on PKD renal epithelial cells may facilitate activation of certain signaling pathways^18^. Consistent with this, renal epithelial primary cilia were either ablated or shortened in adult *Pkd; Ift-B* dko mice, suggesting that shortening or ablating primary cilia may result in inhibition of a pro-cystogenic pathway^16,17^. Yet unlike IFT-B genes, deletion of IFT-A gene, *Thm1*, did not suppress hepatic cystogenesis in adult *Pkd2* cko mice. Additionally, although *Thm1* deletion suppressed cilia length in *Pkd2* cko mice, these shortened *Pkd2;Thm1* dko primary cilia did not counter the increased Notch signaling in cyst-lining cholangiocytes. A possibility is that aberrant activation of Notch in *Pkd2* cko cells occurs through a different mechanism than the downregulation of Notch signaling in *Thm1* cko biliary epithelial cells. Elucidating the mechanism of Notch misregulation in *Thm1-* and *Pkd2*-deficient cholangiocytes would be informative and could lead to identification of additional molecular targets. Further, since IFT-B and -A proteins differentially regulate certain signaling pathways, extending analysis of Notch regulation to *Pkd;Ift-B* dko biliary epithelial cells which show attenuated cystogenesis, would also further mechanistic understanding.

In ADPKD, several pathways that are upregulated in cyst-lining renal epithelial cells, including cAMP, EGFR and ERK signaling, have also been shown to be upregulated in cyst-lining cholangiocytes^35-37^. In ADPKD renal cystogenesis, we have shown that cyst-lining renal epithelial cells have increased Notch signaling^38^. Here we observe that Notch signaling is increased in cyst-lining cholangiocytes of both juvenile and adult ADPKD models, suggesting Notch signaling may be an important driver of hepatic cystogenesis as well. Increased Notch signaling is a major signaling pathway driving cholangiocarcinoma^39,40^. In the kidney, certain underlying pathogenic mechanisms of cancer and PKD overlap^41^. Similarly, in the biliary system, aberrant activation of Notch signaling promoting proliferative disease progression may be an overlapping mechanism.

The contrasting effect on hepatic cystogenesis between deletion of *Ift-B* genes and deletion of *Thm1* in adult ADPKD mouse models is likely due to differing roles of IFT-B and –A in regulating signaling pathways in the liver. Our data also reveal functional differences for *Thm1* between biliary and renal epithelial cells. Perinatal loss of *Thm1* alone does not cause liver cysts, and together with loss of *Pkd2*, does not affect hepatic cystogenesis. Yet in the kidney, perinatal deletion of *Thm1* causes cysts^20^, and deletion of *Thm1* in juvenile *Pkd2* cko mice affects cystogenesis in a tubular-dependent manner. Further, deletion of *Thm1* in adult *Pkd2* cko mice attenuates renal cystogenesis^42^. Thus, *Thm1* influences cystogenesis in the kidney but not in the liver. Elucidating the differences between the roles of *Thm1* in biliary versus renal epithelial cells may help to identify key protein(s) that promote cyst formation or progression. In addition to cell proliferation, fluid secretion is an essential component of cyst growth^43^, and perhaps *Thm1* directly or indirectly regulates a protein that is involved in fluid secretion in the kidney but not in the liver. In summary, our study is the first to demonstrate a role for IFT-A deficiency in liver-associated ciliopathies and provides a new genetic mouse model to study biliary development. Importantly, our data are also the first to show misregulation of Notch signaling in an IFT mouse model of biliary ciliopathy and in PLD, revealing a pathway that warrants further exploration as a novel potential therapeutic target for both diseases.

## Acknowledgments

We thank members of the Jared Grantham Kidney Institute for helpful discussions. We also thank Jing Huang of the KUMC Histology Core and acknowledge support of this core (Intellectual and Developmental Disabilities Research Center NIH U54HD090216; COBRE NIH P30GM122731). This work was also supported by a K-INBRE Summer Student Award to JTC (K-INBRE P20GM103418), R01DK108433 to MS, and R01DK103033 to PVT.

## Contributions

WW, TSP, HHW, RD, MAK, JTC, MTP and PVT performed experiments. WW, TSP, HHW, RD, MAK, JTC, UA, MTP, MS and PVT analyzed and interpreted data. WW, UA, MTP, MS and PVT designed research. WW and PVT wrote the manuscript.

## Supplementary Figure Legends

**Figure S1.**
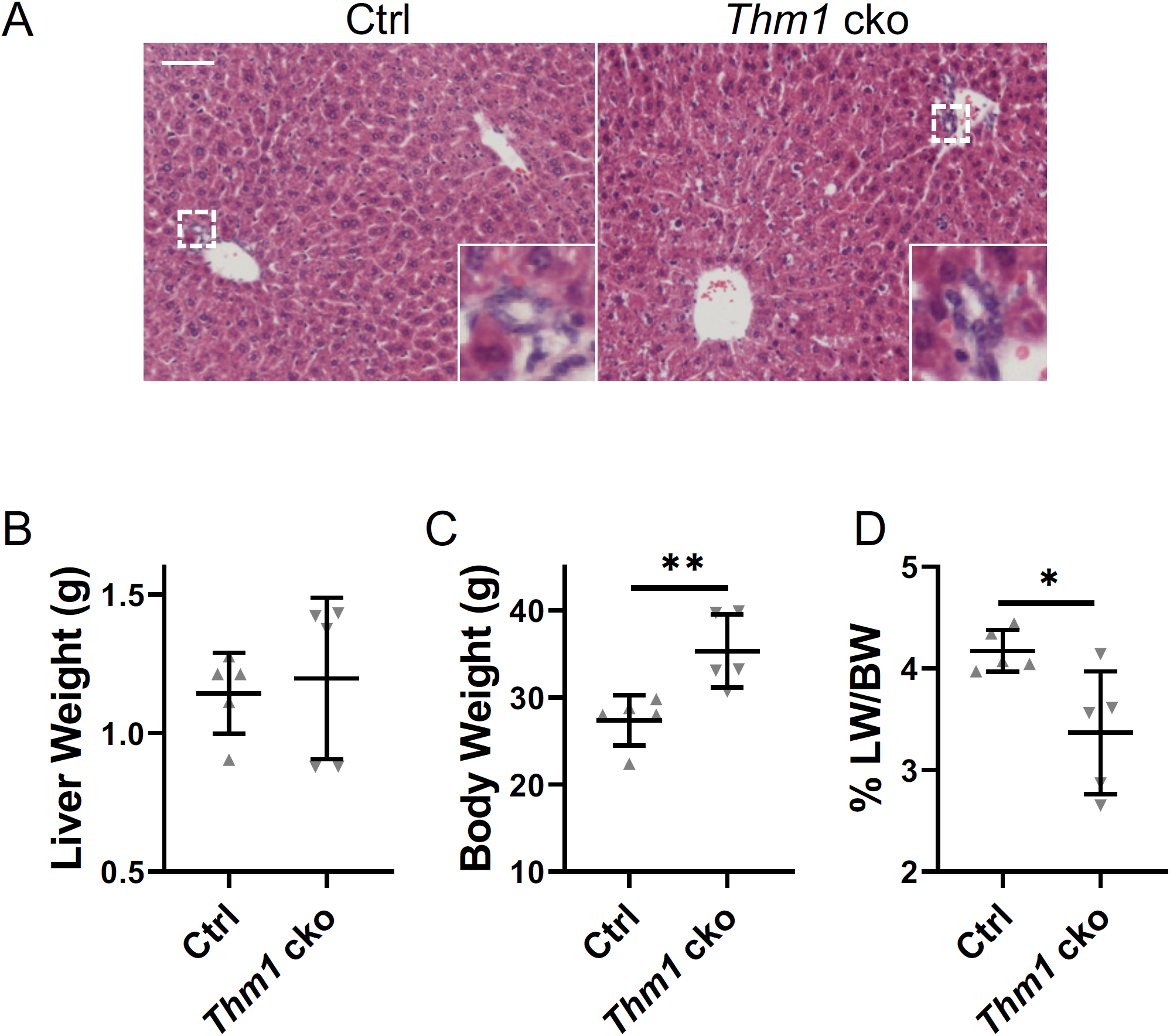
Liver histology of adult-onset *Thm1* cko mice. (A) H&E-stained liver sections of 6-month old mice. Scale bar -50µm (B) Liver weight (C) Body weight (D) Liver weight/Body weight ratios. Data points represent individual mice. Statistical significance was determined by unpaired t-test in (C) and (D). *P<0.05; **P<0.01

**Figure S2.**
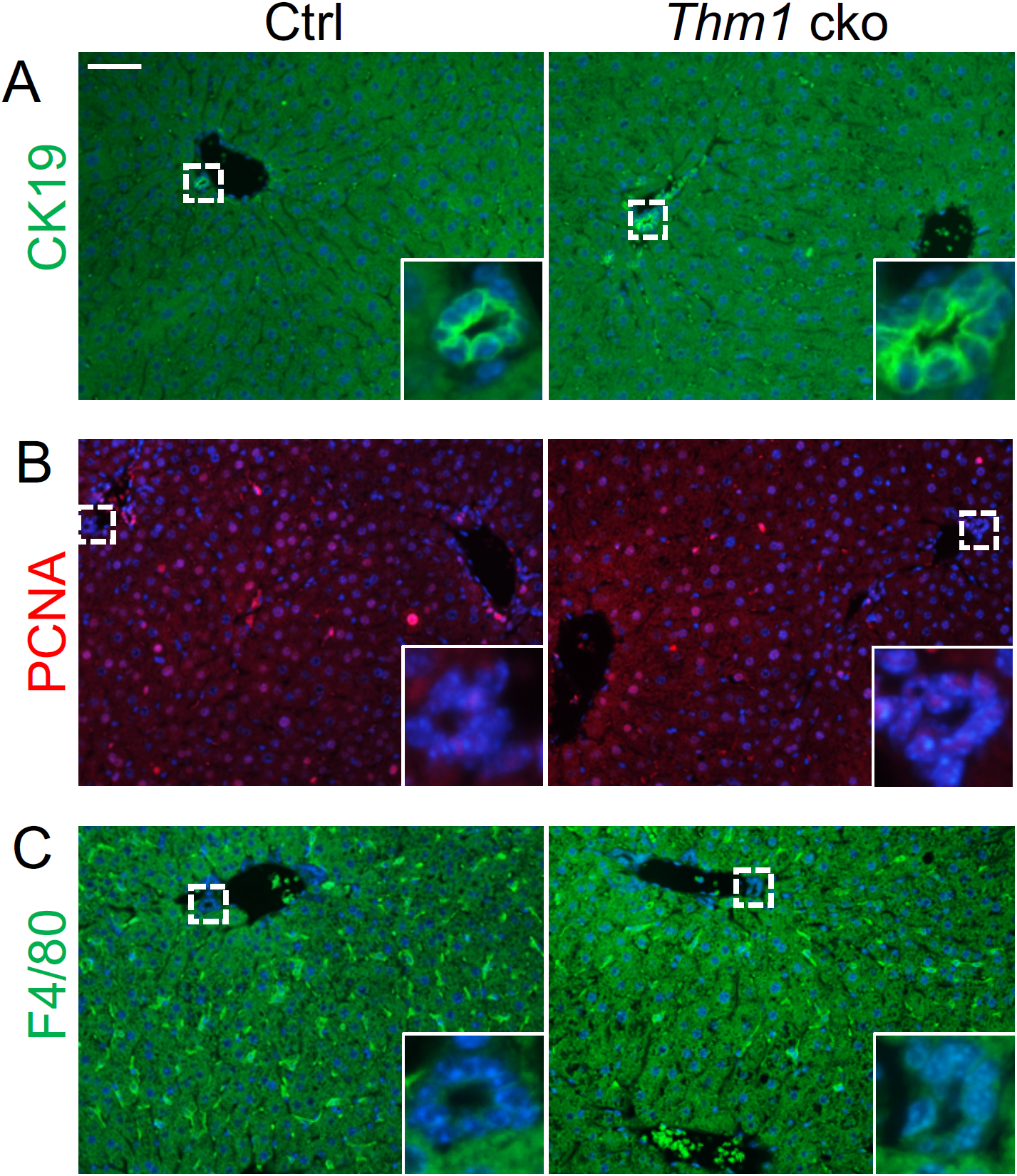
Biliary ducts of adult onset *Thm1* cko livers resemble control. Immunostaining for (A) CK19 (green); (B) PCNA (red) and; (C) F4/80 (green). N=3 mice/genotype

**Figure S3.**
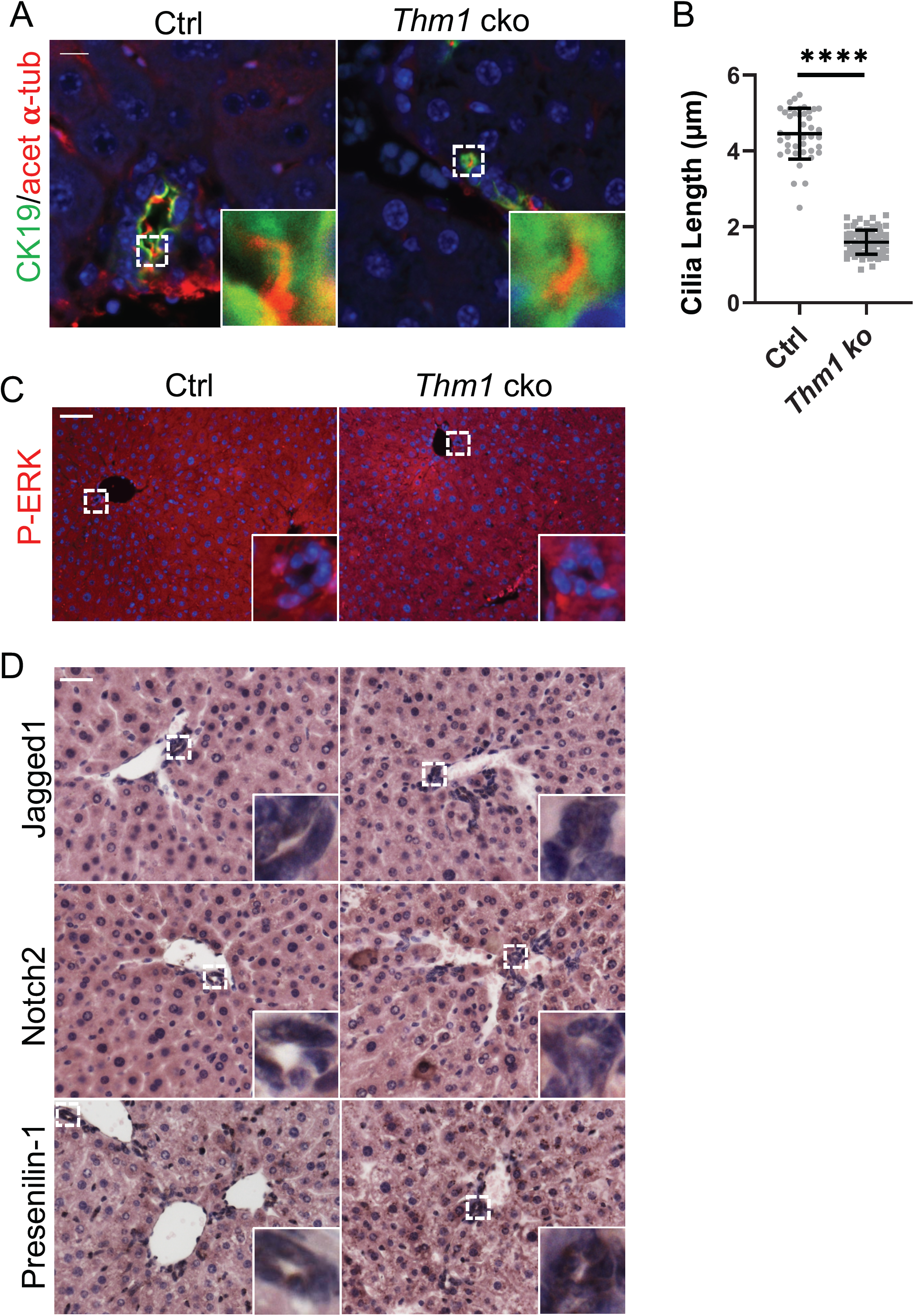
Primary cilia and ERK and Notch signaling in biliary regions of adult *Thm1* cko mice. (A) Immunostaining for acetylated α-tubulin (red) and CK19 (green) of P21 mice. Scale bar -10 µm. (B) Quantification of cilia lengths. Each data point represents an individual cilium from 3 mice/genotype. Statistical significance was determined by unpaired t-test. ****P< 0.0001. (C) Immunostaining for P-ERK. (D) Immunohistochemistry for Notch signaling components – Jagged1 ligand, Notch2 receptor and Presenilin-1. Dotted boxed regions are shown at higher magnification in insets. N=3 mice/genotype. Scale bar -50 µm

